# Species-level resolution of 16S rRNA gene amplicons sequenced through the MinION^TM^ portable nanopore sequencer

**DOI:** 10.1101/021758

**Authors:** Alfonso Benítez-Páez, Kevin J. Portune, Yolanda Sanz

**Affiliations:** Microbial Ecology, Nutrition & Health Research Unit. Institute of Agrochemistry and Food Technology Institute, National Research Council (IATA-CSIC), Valencia, Spain.

**Keywords:** MinION, Nanopore sequencer, 16S rDNA amplicon sequencing, Microbial diversity, Long-read sequencing

## Abstract

**Background:** The miniaturised and portable DNA sequencer MinION^TM^ has been released to the scientific community within the framework of an early access programme to evaluate its application for a wide variety of genetic approaches. This technology has demonstrated great potential, especially in genome-wide analyses. In this study, we tested the ability of the MinION^TM^ system to perform amplicon sequencing in order to design new approaches to study microbial diversity using nearly full-length 16S rDNA sequences.

**Results:** Using R7.3 chemistry, we generated more than 3.8 million events (nt) during a single sequencing run. These data were sufficient to reconstruct more than 90% of the 16S rRNA gene sequences for 20 different species present in a mock reference community. After read mapping and 16S rRNA gene assembly, consensus sequences and 2d reads were recovered to assign taxonomic classification down to the species level. Additionally, we were able to measure the relative abundance of all the species present in a mock community and detected a biased species distribution originating from the PCR reaction using ‘universal’ primers.

**Conclusions:** Although nanopore-based sequencing produces reads with lower per-base accuracy compared with other platforms, the MinION^TM^ DNA sequencer is valuable for both high taxonomic resolution and microbial diversity analysis. Improvements in nanopore chemistry, such as minimising base-calling errors and the nucleotide bias reported here for 16S amplicon sequencing, will further deliver more reliable information that is useful for the specific detection of microbial species and strains in complex ecosystems.

## Background

The third generation of DNA sequencers is based on single-molecule analysis technology that constantly is under development to minimise errors and produce high quality reads. Oxford Nanopore Technologies (ONT) released the first miniaturised and portable DNA sequencer to researchers in early 2014, within the framework of the MinION^TM^ Access Programme. The MinION^TM^ is a USB stick-sized device operated from a computer via USB 3.0. Real-time data analysis can be visualised in terms of number of reads and length distribution. Nucleotide base-calling and quality assessment of reads require further processing, where data exchange of Hierarchical Data Format (HDF5) files, containing a large amount of numerical data, is indispensable. This data exchange is done via the Internet through the Metrichor platform; a process that can optionally be launched after the sequencing process itself. According to its theoretical capabilities, the MinION^TM^ provides new alternatives for genomic analyses. One of the most attractive capabilities of the MinION^TM^ platform is the sequencing of complete bacterial genomes, as demonstrated recently by Quick et al. [1]. Another major advantage of the MinION^TM^ platform, compared to other popular sequencing technologies, is its performance in terms of read length. Theoretically, nanopore-sensing technology is able to generate thousands of reads that are hundreds to thousands of nucleotides in length; the only limitation being the DNA fragments generated during nucleic acid extraction procedures, which frequently produce fragmented DNA with an average length of 50kb. Although short-read length sequencing approaches deliver high quality sequences, these partial genome sequences with unsolved repetitive elements make it impossible to study genetic variation or molecular evolution directly or indirectly associated to such elements. Therefore, long-read approaches offer new insights into genomic analysis, facilitating the assembly of complete genomes through hybrid strategies [2]. In addition to genome sequencing analysis, microbial diversity and taxonomic approaches are also deeply limited by short-read strategies. Early massive sequencing approaches producing 50 nt (Genome Analyzer, Solexa/Illumina) to 200 nt (454 Roche) effective reads with a modest average quality only allowed accurate exploration of diversity at the phylum level. However, thanks to improvements in the chemistry of the most common, popular sequencing platforms in recent years, it is now possible to characterise microbial communities in detail down to the family or even genus level. To date, paired-end short read approaches for massive sequencing permit the analysis of sequence information of roughly 30% (∼500nt) of the full 16S rRNA gene, which means taxonomic assignment of reads at the species level is elusive. Therefore, implementation of long-read sequencing approaches to study 16S rRNA genes will permit the design of new studies to provide evidence for the central role of precise bacterial species/strains in a great variety of microbial consortia. As a consequence, we present a preliminary study of 16S rDNA amplicon sequencing of a mock microbial community composed of genomic DNA from 20 different bacterial species (BEI Resources) using the MinION^TM^ sequencing platform. The aim of this study is to evaluate the application of nanopore technology in performing bacterial diversity and taxonomic analysis on nearly full-length bacterial 16S rRNA genes.

## Data description

Raw data collected in this experiment were obtained as fast5 files using MinKNOW software v0.50.1.15 (Oxford Nanopore Technologies), after conversion of electric signals into base calls via the Metrichor Agent v2.29 and the 2D Basecalling workflow v1.16. Base-called data passing quality control and filtering were downloaded and basic statistical analysis was carried out using *poretools* [3] and *poRe* [4]. Mapping statistics are depicted in Table 1. Fast5 raw data can be accessed at the European Nucleotide Archive (ENA) under the project ID PRJEB8730 (sample ERS760633). Only one data set was generated after a sequencing run of MinION^TM^.

## Analyses

DNA reads derived from MinION sequencing can be classified into three types: ‘template’, ‘complement’, and ‘2d’ reads. While template reads come from DNA strands that are primed by a leader adapter and passed through the pore, the complement reads are generated only if a second adapter (hairpin adapter) is present in the same DNA fragment, thus permitting sequencing both strands of a single molecule in a concatenated manner. The 2d reads are products of aligning and merging sequences from template and complement reads generated from the same DNA fragment: these contain a lower error rate, owing to strand comparison and mismatch correction. After the sequencing process, we obtained 3,404 reads, of which 58.5% were template reads (1,991), 23.8% were complement reads (812), and 17.7% were 2d reads (601). Read lengths had a wide distribution ranging from 12 nt to more than 50,000 nt in length, with a median of 1,100 nt. We hypothesised that extremely large reads might be products of the ligation of multiple amplicons. However, when we tried to align these large reads to reference sequences, we detected no matches (data not shown). Accordingly, a filtering step was performed by retaining 97% of the original dataset (3,297 reads), with a size range between 100 and 2,000nt in length for downstream analysis.

In the first step of our analysis, we used the large set of template and complement reads to assess the global performance of the amplicon sequencing process. Consequently, we analysed basic read-mapping statistics to uncover potential pitfalls of the MinION platform and tried to reconstruct the reference sequences. We assembled more than 90% of 16S rRNA gene sequences for all organisms included in the mock community (Table1). We observed that even at very low coverage, such as that retrieved for the *Bacteroides vulgatus* 16S rRNA gene (Figure 1), it is possible to reconstruct almost 93% of the entire gene. Indeed, the maximum size of amplicons sequenced in all cases was close to the expected amplicon size according to the universal primers used in the PCR design (Table 1). In terms of coverage, we hypothesised that the lower than expected number of 16S reads from *B. vulgatus* species (Figure 1) resulted from a bias caused during PCR amplification, despite using high coverage primers [5, 6], or as a result of the sequencing process itself. To further investigate this matter, we performed an absolute quantification of 16S rRNA genes using qPCR from three different species with a high coverage, close-to-expected coverage, and the lowest coverage, respectively, which were present in the initial PCR sample used for library construction and sequencing. A correlation between the number of molecules present in the starting material and the coverage obtained after the sequencing process (Pearson’s r = 0.99, p≤0.0514) was detected (Figure 2), indicating that the sequencing process faithfully reproduced the proportion of amplicons present in the sample and the coverage bias was therefore derived from the starting material generated by PCR. Despite this bias, the 16S rRNA gene from *B. vulgatus* was fully assembled with low variation (25/1,403 = 1.78%) after DNA read alignment and pileup (Table 1).

**Figure 1.**
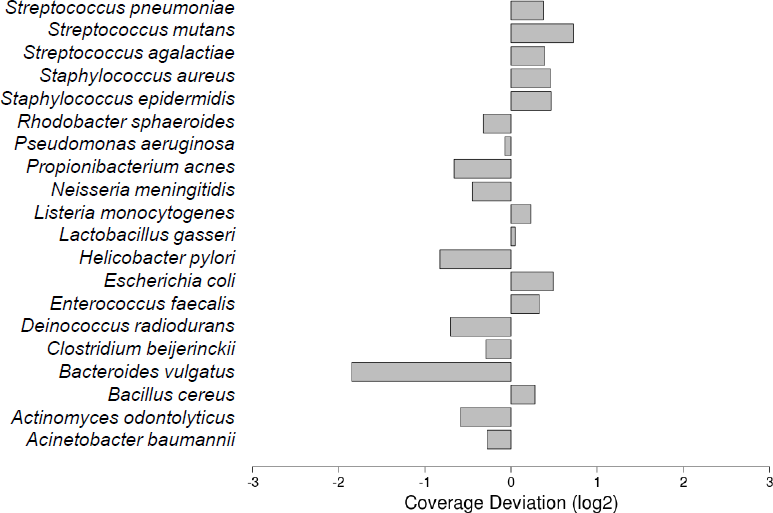
Species abundance in the mock community detected by MinION^TM^. Species coverage was calculated by obtaining the fold-change (Log_2_) of species-specific read counting against the expected average for the entire community. A coverage bias was assumed when coverage deviation was lower than -1 or higher than 1.

**Figure 2.**
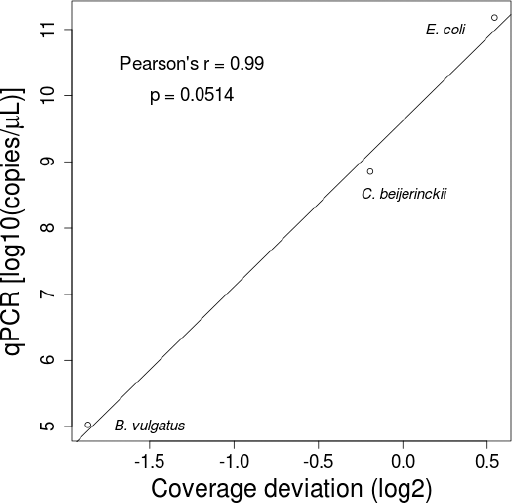
Sequencing coverage versus copies of respective 16S rRNA genes present in the starting material. Scatter plot of the coverage deviation calculated for *B. vulgatus, C. beijerinkii,* and *E. coli* against the calculated number of respective 16S copies present in the PCR sample used as starting material of the sequencing reaction.

Read-mapping statistics were analysed to further measure the performance of MinION^TM^ sequencing in microbial diversity analysis based on 16S rDNA sequences. The GC content of reads produced by MinION^TM^ showed an important and significant correlation (Pearson’s r = 0.47, p≤0.0376) against the GC content of reference values (Figure 3A), which indicates that the GC content of 16S rDNA sequences is fairly well replicated during sequencing. However, we found a 16S rDNA GC content bias, to some extent, in the reads obtained from MinION^TM^, which inalmost all cases exceeds the GC content of the reference (Figure 3A). To test the probable influence of GC content bias in base-calling accuracy, we performed linear comparisons against mismatch rates, indel rates, and coverage deviation. We observed that both coverage deviation (p≤0.00003) and mismatch rate (p≤0.00004) are significantly influenced by read GC content (Figure 3B and 3C, respectively). In the first case, the influence of GC content on coverage deviation could have a minimal effect because 95% of species analysed show no more than a one-fold deviation. However, with GC bias detected in reads from the MinION^TM^ sequencer, this effect could be magnified, especially in species where GC content is high. On the other hand, we found a strong correlation between the GC content of reads and the mismatch rate retrieved from alignments, which would insinuate again that GC content is a factor that influences 16S rDNA amplicon sequencing in the MinION^TM^ platform. Conversely, GC content did not appear to profoundly affect indel rate (Figure 3D).

**Figure 3.**
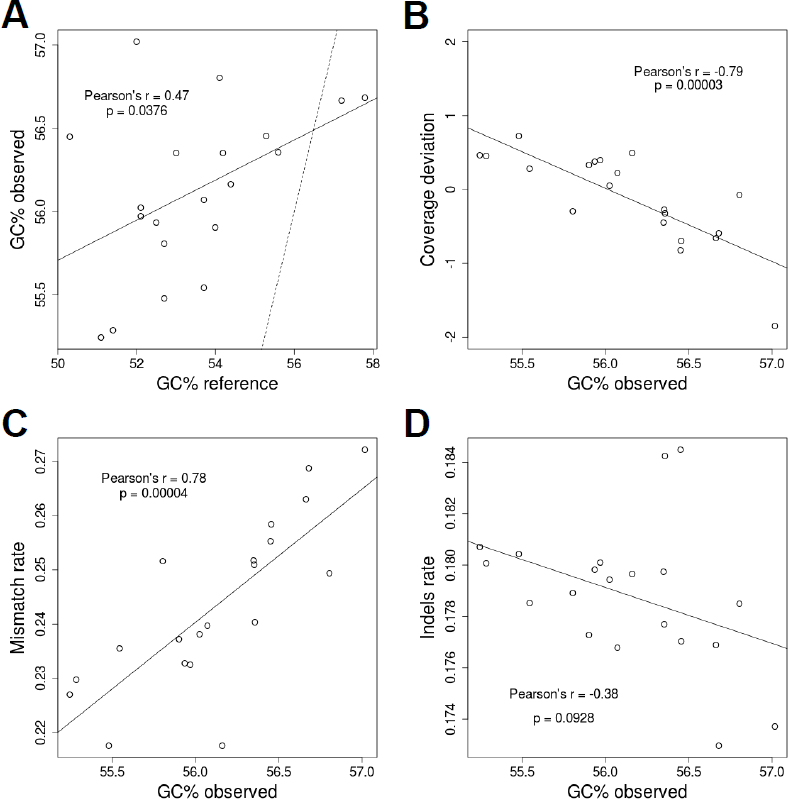
Per-base accuracy of the mapped reads. (A)Scatter plot of the GC content observed in mapped reads against the GC obtained from the reference sequences. The dashed line indicates a correlation with Pearson’s r = 1. (B) Correlation between the GC content observed in mapped reads and coverage bias observed in Figure 1. (C) Influence of the GC content observed in mapped reads on mismatch rates calculated after mapping. (D) Scatter plot of the observed GC content of mapped reads and indel rates calculated after mapping. In all cases the Pearson’s r coefficients and p values supporting such correlations are presented inside the scatter plots and solid lines indicate the tendency of correlations.

The complete assembly of the amplified 16S rRNA gene permitted the quantification of the level of sequence variants in the consensus sequence. These variants were recovered after a pileup of reads against reference sequences, and they were variable in number with a median of 8 variants per 16S rRNA gene (Table 1). This number of nucleotide substitutions means that approximately 0.5% of the 16S rDNA sequence assembled from MinION^TM^ reads retained unnatural genetic variants directly generated from the sequencing process itself, theoretically allowing a bona fide identification and taxonomic assignment of 16S rDNA sequences at the species level. In the worst cases, where the number of variants were meaningful (∼2.3% of the full assembly), such as those observed for *Acinetobacter baumannii* and *Bacillus cereus* (Table 1), direct BLAST comparisons of these assembled 16S rDNA sequences against the NCBI 16S database only produced matches with homologous sequences belonging to the same species, respectively (data not shown).

A final step in our analysis tested whether or not the information obtained through sequencing of nearly full-size 16S rRNA genes using the MinION^TM^ platform is useful to perform taxonomic assignment with tools commonly employed in microbial community analysis. For this aim, we used the information derived from 2d reads (601 reads), which is limited in terms of the effective number of reads but more reliable in terms of sequence identity. Using the SINA web service [7] we obtained the taxonomic assignment of 2d reads to the Silva bacterial 16S database [8]. The results of this approach are shown in Table 2. Out of the 17 different genera present in the mock community, we retrieved information for six of them, with an assignment threshold of 80%, seven with 70% and eight with 60%. Using 60% as the lowest assignment threshold, we started to retrieve unexpected genera composition in our 2d data set, indicating that reliable identifications must be set with a higher identity threshold. As expected, taxonomy assignation was limited to those species with a higher coverage during sequencing processing (Figure 1), which is consistent with the number of 2d reads expected after aligning and merging respective template and complement reads obtained from the 16S rRNA genes of species over-represented in the starting material. We expect that the whole repertoire of species present in the sample can be detected by increasing the performance of the sequencing process. This would allow us to obtain a larger raw dataset and, particularly, more 2d reads containing more reliable information to perform taxonomic assignments and disclose the full inventory of species present in the microbial community under study. Finally, a BLAST-based assignment against the NCBI bacterial 16S rRNA gene database retrieved the identities of 8 of the 20 species presented in the mock community analysed (Table 2). Although other species were also retrieved, they exhibited a high level of affiliation in terms of the 16S rDNA sequence identity to the true species included in the mock community.

## Discussion

The inventory of microbial species based on 16S rDNA sequencing is frequently used in biomedical research to determine microbial organisms inhabiting the human body and their relationship with disease. Identification of microbial species inhabiting different areas and cavities of the human body currently relies on the handling and processing of millions of DNA sequences obtained through the second generation of massive and parallel sequencing methods. However, these methods are still limited, mainly in terms of DNA read length. The inability to determine complete 16S rDNA sequences during massive sequencing has led to the development of multiple algorithms dedicated to theoretically discerning microbial species present in samples according to the sequence similarity degree, or Operational Taxonomic Units (OTUs). Despite high accuracy and a constant update of the methods used in OTU-based approaches, available algorithms produce no consensus outputs, leaving a high degree of uncertainty when the number of theoretical species and their abundance is the subject of study [9-12].

Thanks to the fact that they overcome DNA read limitations at the expense of decreasing throughput, a third generation of sequencing methods based on single-molecule technology offers new possibilities to study microbial diversity and taxonomic composition. MinION^TM^ is one of these single-molecule methodologies, which has demonstrated its capacity in genome sequencing [1, 13]. Recent studies have reported the application of this technology in medical microbiology by using amplicon sequencing to determine bacterial and viral infections [14, 15]. Our results indicate that the MinION^TM^ per-base accuracy (65–70% for template reads, and 85% for 2d reads) is in concordance with previous results [1, 14, 16]. We found that sequence coverage was close to expected values in most cases, with the exception of that of *B. vulgatus* (gene GC = 52%), which was 1.84-fold less than the expected coverage. Using absolute quantification of molecules presented in the starting material, we demonstrated that such coverage bias came from the PCR process used to generate the 16S amplicons, despite using ‘universal’ primers with higher coverage among bacterial species [5]. Despite this, such coverage was enough to reconstruct 93% of the 16S rRNA gene of *B. vulgatus* with a low proportion of unnatural variants.

We observed a general influence of GC content in the mismatch rate but not in the indel rate. This suggests that base miscalling could be associated with the amplicon GC content. Moreover, a slight correlation between the amplicon GC content observed and coverage bias was evidenced, indicating that GC content could be negatively affecting amplicon coverage to some extent. Although MinION^TM^ was able to replicate the GC content expected for every amplicon sequenced fairly well, we observed a slight over-representation of GC in all reads obtained. This over-calling of GC bases in 16S rDNA amplicons could additionally influence the issues stated above in a negative manner. The R7.3 chemistry used in MinION^TM^ allowed the acquisition of reads of moderate quality, which were enough to reconstruct more than 90% of the 16S rRNA gene in all 20 bacterial species analysed. None of the 20 16S rDNA consensus sequences assembled showed more than 3% of sequence variation, which can be considered as a threshold for canonical species identification. Therefore, the consensus sequence assembled was useful to obtain a reliable taxonomic identification at the species level. As expected, unnatural variants were associated with low coverage regions. Therefore, increasing the sequencing coverage will drastically reduce the ambiguities of the assembled sequences. When we tested the high quality reads (2d) in common routines for the analysis of microbial communities, the SINA web server retrieved a taxonomic assignation, indicating the presence of 7 genera out of the 17 expected for the mock community without any mismatches (using 70% sequence identity as a threshold). Although this number of matches can be considered low, it was directly associated with the sequencing coverage, therefore, a larger 2d data set generated from a greater sequencing effort would produce enough information to identify the entire community. In terms of the study of microbial communities, results obtained using 16S rDNA amplicon sequencing through the MinION^TM^ device are promising. Despite the observed modest per-base accuracy of this sequencing platform, we were able to reconstruct nearly full-length16S rDNA sequences for 20 different species analysed from a mock bacterial community, and were able to obtain an acceptable taxonomy assignation for high quality 2d reads, only limited by the sequencing effort. This seems to be the major handicap of the MinION^TM^ platform for microbial diversity analysis. To date, MinION^TM^ and nanopore technologies have demonstrated great potential in DNA sequencing by allowing the retrieval of whole bacterial genome sequences with a minimum level of variation [1]. With the results presented here, we postulate that the MinION^TM^ platform is a reliable methodology to study the diversity of microbial communities. It permits: i) a taxonomic identification at the species level through 16S rDNA sequence comparisons, and ii) a relative quantification to determine the species abundance. This type of analysis will likely become more accurate over time as nanopore chemistry is improved in future releases, with the concomitant increasing of the throughput, pivotal to disclose the hundreds of species present in complex microbial communities. The implementation of the “What’s In My Pot” (WIMP) Metrichor workflow, which aims to acquire real-time taxonomic sequence identification by comparing against different bacterial references databases (i.e. NCBI, SILVA [8], GreenGenes [17]), will be helpful in other types of analyses related to those presented here. Accordingly, sequence studies of the entire 16S rDNA molecule could allow OTU-based analysis to be bypassed completely, thus making it feasible to obtain a direct inventory of bacterial species and relative abundance, as well as to determine the key players at the species level in different microbial communities of interest.

## Methods

### Bacterial DNA and 16S rDNA amplicons

Genomic DNA for the reference mock microbial community was kindly donated by BEI Resources [18]. This mock community (HM-782D) is composed of a genomic DNA mix from 20 bacterial strains containing equimolar ribosomal RNA operon counts (100,000 copies per organism per μL), as indicated by the manufacturer. According to instructions provided by BEI Resources, 1 μL of mock community DNA was used to amplify 16S rRNA genes. DNA was amplified by 30 PCR cycles at 95°C for 20 s, 47°C for 30 s, and 72°C for 60 s. Phusion High-Fidelity Taq Polymerase (Thermo Scientific) and the primersS-D-Bact-0008-c-S-20 and S-D-Bact-1391-a-A-17, which target a wide range of bacterial 16S rRNA genes, were used during PCR [5, 6]. Amplicons consisted of∼1.5kbp blunt-end fragments, which were purified using the Illustra GFX PCR DNA and Gel Band Purification Kit (GE Healthcare). Amplicon DNA was quantified using a Qubit 3.0 fluorometer (Life Technologies).

### Amplicon DNA library preparation

The Genomic DNA Sequencing Kit SQK-MAP-005 was used to prepare the amplicon library to be loaded into the MinION^TM^. Approximately 250 ng of amplicon DNA (0.25 pmol) was processed for end repair using the NEBNext End Repair Module (New England Biolabs), followed by purification using Agencourt AMPure XP beads (Beckman Coulter). Subsequently, according to the manufacturer’s instructions, we used 200 ng of the purified amplicon DNA (∼0.2 pmol) to perform dA-tailing using the NEBNext dA-tailing module (New England Biolabs) with a total volume of 30 μL, and incubated the sample at 37°C for 15 minutes. Fifty μl of Blunt/TA ligase master mix (New England Biolabs), 10 μL of adapter mix, and 2 μL of HP adapter were added to the 30 μl dA-tailed amplicon DNA. The reaction was incubated at 16°C for 15 minutes. The adapter-ligated amplicon was recovered using Dynabeads^®^ His-Tag (Life Technologies) and washed with washing buffer provided with the Genomic DNA Sequencing Kit SQK-MAP-005 (Oxford Nanopore Technologies). Finally, the sample was eluted from the Dynabeads^®^ by adding 25 μL of elution buffer and incubating for 10 minutes at room temperature before pelleting in a magnetic rack.

### Flowcell set-up

A brand new, sealed R7.3 flowcell was stored at 4°C before first use. It was fitted to the MinION^TM^ with plastic screws, ensuring a good thermal contact. The R7.3 flowcell was primed twice using 71 μL premixed nuclease-free water, 75 μ1 2× running buffer, and 4μLfuel mix. At least 10 minutes were needed to equilibrate the flowcell before each round of priming and before final DNA library loading.

### Amplicon DNA sequencing

The sequencing mix was prepared with 63 μl nuclease-free water, 75 μ1 2× running buffer, 8 μL DNA library, and 4 μL fuel mix. A standard 48-hour sequencing protocol was initiated using the MinKNOW^TM^ v0.50.1.15. Base-calling was performed through data transference using the Metrichor^TM^ agent v2.29.1 and 2D base-calling workflow v1.16. During the sequencing run, one additional freshly diluted aliquot of DNA library was loaded after 12 hours of initial input.

### Data analysis

Quality assessment of read data and conversion to fasta format was performed using the *poretools* [1] and *poRe* [4] packages. Fasta sequences were filtered by retaining those with a length between 100 and 2000 nt. Read-mapping was performed against the 16S ribosomal RNA sequences for the species present in the mock community (see Availability of supporting data).

Read-mapping was performed using the LAST aligner v.189 [19] with parameters -q1 -b1 -Q0 -a1 -e45, which were configured to give the best balance between 16S rDNA assembly length and variants. LAST outputs were converted to sam files and processed with *samtools* [20] to build indexed bam files and obtain consensus sequences from alignments and variant calling. Read-mapping stats from sam files were calculated with the *ea-utils* package and its *sam-stats* function [21]. Different comparisons, GC content correlations, and plots were performed and drawn in R v3.2.0 [22]. Species coverage was calculated by obtaining fold-change (Log_2_) of species-specific read counting against the expected average for the entire community. A coverage bias was assumed when coverage deviation was lower than -1 or higher than 1. Taxonomy assignment of 2d reads was performed using the Silva database [8] and the SINA aligner [20]. Sequences were submitted to the SINA alignment web server using 80%, 70%, and 60% of identity thresholds to ensure a reliable identification. Additional identification at the species level was done using BLAST and the reference NCBI 16S rDNA database.

For the absolute quantification of 16S amplicons we used the following primers: *Escherichia coli* GGACGGGTGAGTAATGTCTGG and ACCTACTAGCTAATCCCATCTG; *Clostridium beijerinkii* AGAACCTTACCTAGACTTGACATC and GCTACTAACAATAAGGGTTGCG; and *Bacteroides vulgatus* CACGGGTGAGTAACACGTATCC and GCATCCCCATCGTCTACCGGAA. Single-stranded DNA (ssDNA), fully covering the respective 16S rDNA regions to amplify for the *E. coli, C. beijerinckii, and B. vulgatus* species, was obtained from Isogen Life Science B.V (Utrecht, The Netherlands) where it was synthesized, PAGE-purified, and quantified and used in molecule titration for qPCR. The qPCR was performed on a LightCycler^®^ 480 instrument (Roche Life Science) using the SYBR Green I Master Mix reagent (Roche Life Science), 0.625μM oligos, and 1 μL of 1:20 diluted and purified PCR product generated with ‘universal primers’. After 35 cycles of amplificationat 95°C for 10 s, 64°C for 20 s, and 72°C for 15 s, absolute quantification was determined using LightCycler^®^ 480 SW v1.5 software (Roche Life Science). Ct values were obtained from serial dilutions of respective ssDNA with known concentrations.

## Availability of supporting data

Accessions for the 16S ribosomal RNA sequences for the species present in the mock community are available at GenBank: NC_009085 range c3505652-3504124, NZ_GG753639 range 96928-98455, NC_003909 range82453-83960, NC_009614 range c4744649-4743140, NC_009617 range c5775228-5773724, NC_001263 range c2287019-2285518, NC_017316 range 213429-214977, NC_000913 range 4208147_4209688, NC_000915 range c1512634-1511137, NC_008530 range c1560731-1559153, NC_003210 range 243556-245041, NC_003112 range c2137452-2415909, NC_006085 range 606163-607687, NC_002516 range c6044743-6043208, NC_007493 range 1-1463, NC_010079 range c2003413-2001874, NC_004461 range c1816154-1811601, NC_004116 range 348575-350125, NC_004350 range 185749_187300, and NC_003028 range c1815064-1813505). Alternatively, a multi-fasta file containing the 16S reference sequences for the species included in the mock community is available at https://github.com/alfbenpa/16S_MinION.

Further supporting data can be found in the *GigaScience* database, GigaDB [23].

## Abbreviations

BLAST: Basic Local Aligment Tool
EC: European Commission
ENA: European Nucleotide Archive
HDF: Hierarchical Data Format
NCBI: National Center for Biotechnology Information
ONT: Oxford Nanopore Technologies
OTU: Operational Taxonomic Unit
PCR: Polymerase Chain Reaction
rDNA: DNA encoding for the Ribosomal RNA
rRNA: Ribosomal RNA
SINA: SILVA Incremental Aligner
USB: Universal Serial Bus
WIMP: What’s In My Pot Metrichor Workflow.

## Acknowledgements

Authors thank the European 7th Framework Programme for funding ABP and KP, who were supported by the EC Project no. 613979 (MyNewGut).

## Authors’ contributions

ABP and YS designed the study and managed the project. ABP performed the experiments, and analysed and managed the data. ABP, KP, and YS wrote the manuscript. All authors read and approved the final manuscript.

